# The impact of water deficit and heat stress combination on the molecular response, physiology and seed production of soybean

**DOI:** 10.1101/2020.09.30.320341

**Authors:** Itay Cohen, Sara I. Zandalinas, Felix B. Fritschi, Soham Sengupta, Yosef Fichman, Rajeev K. Azad, Ron Mittler

**Author notes:** Contributed equally (Itay Cohen, Sara I. Zandalinas, Felix B. Fritschi). **Author emails:** I.C.,; S.I.Z.,; F.B.F.,; S.S.,; Y.F., R.K.A,; R.M. **Abbreviations:** Control, CT; water deficit, WD; heat stress, HS; combined water deficit and heat stress, WD + HS; relative water content, RWC; fresh weight, FW; turgid weight, TW; dry weight, DW; transcription factor, TF; heat shock factor, HSF; Dehydration responsive element binding, DREB; Plant Introduction, PI.

## Abstract

A combination of drought and heat stress, occurring at the vegetative or reproductive growth phase of many different crops, can have a devastating impact on yield. In soybean (*Glycine max*), a considerable effort has been made to develop genotypes with enhanced yield production under conditions of drought or heat stress. However, how these genotypes perform in terms of growth, physiological responses and most importantly seed production, under conditions of drought and heat combination is mostly unknown. Here, we studied the impact of water deficit and heat stress combination on the physiology, seed production and yield per plant of two soybean genotypes, Magellan and Plant Introduction (PI) 548313, that differ in their reproductive responses to heat stress. Our findings reveal that although PI 548313 produced more seeds than Magellan under conditions of heat stress, under conditions of water deficit and heat stress combination its seed production decreased. Because number of flowers and pollen germination of PI 548313 remained high under heat or water deficit and heat combination, the reduced seed production exhibited by PI 548313 under the stress combination could be a result of processes that occur at the stigma, ovaries and/or other parts of the flower following pollen germination.

**Highlight:** Tolerance to heat stress was found not to confer tolerance to a combination of water deficit and heat stress in soybean, highlighting the need for breeding strategies targeting the stress combination.

## Introduction

Heat waves occurring during periods of prolonged drought stress can have a devastating impact on plant growth, agricultural productivity and yield (Mittler and Blumwald, 2010; Suzuki *et al*., 2014; Choudhury *et al*., 2017). A combination of drought and heat stress occurring during the summers of 1980, 1988, 2000, 2003 and 2008 in the US, for example, resulted in yield losses estimated in 33.3, 44.8, 7.6, and 7.1 billions of US dollars, respectively (Mittler, 2006; https://www.ncdc.noaa.gov/billions/events). Similar incidents occurring worldwide have been the cause of economic and social instability, as well as mass migrations (*e.g*., Missirian and Schlenker, 2017; Ault, 2020). Multiple studies have shown that the impact of drought and heat combination on the growth, yield, harvest index, and seed quality of different plants and crops is significantly higher compared to that of drought or heat stress applied individually (Cohen *et al*., 2020). Moreover, the molecular, physiological, metabolic and proteomic responses of plants to a combination of drought and heat stress was shown to be unique compared to that of drought or heat stress applied alone (*e.g*., Rizhsky *et al*., 2004; Mittler, 2006; Rasmussen *et al*., 2013; Prasch and Sonnewald, 2013; Suzuki *et al*., 2014; Shaar-Moshe *et al*., 2017, 2019). Among the many unique physiological features of a combination of drought and heat stress is the conflicting stomatal response of plants to this stress combination (Rizhsky *et al*., 2004; Mittler, 2006; Balfagón *et al*., 2020; Zandalinas *et al*., 2020*b*). In response to heat stress plants open their stomata to cool their leaves via transpiration, while in response to drought plants close their stomata to avoid water loss. During a combination of drought and heat stress, however, the outcome of this conflict, at least in Arabidopsis and tobacco, is that stomata remain close (Rizhsky *et al*., 2002, 2004). As a result, leaf temperature of Arabidopsis and tobacco plants subjected to a combination of drought and heat stress is significantly higher compared to that of plants subjected to drought or heat stress alone (Rizhsky *et al*., 2002, 2004). In addition to the different physiological differences observed between plants subjected to drought, heat stress or drought and heat stress combination, the state of stress combination was found to be accompanied by altered expression of thousands of transcripts that are unique to it and do not occur in response to drought or heat stress, applied individually to plants (Rizhsky *et al*., 2004; Rasmussen *et al*., 2013; Prasch and Sonnewald, 2013; Shaar-Moshe *et al*., 2017, 2019; Zandalinas *et al*., 2020*b*). Similar observations were also made for many proteins and metabolites that specifically accumulate in plants during stress combination (Rizhsky *et al*., 2004; Koussevitzky *et al*., 2008). Moreover, in a recent genome-wide association study, single nucleotide polymorphism (SNPs) markers were found to be specifically associated with drought and heat stress combination, further highlighting the uniqueness of this type of response (Yuan *et al*., 2019).

Because episodes of drought and heat stress combination are expected to increase in intensity and frequency in the coming years, due to the largely unopposed process of global warming, and its consequential impacts on climate change (*e.g*., Mittler and Blumwald, 2010; Challinor *et al*., 2014; Mazdiyasni and AghaKouchak, 2015; Borghi *et al*., 2019; Ault, 2020; Grossiord *et al*., 2020), breeding crops for enhanced tolerance to this stress combination is of the utmost importance for modern agriculture (Mittler and Blumwald, 2010; Bailey-Serres *et al*., 2019; Zandalinas *et al., 2020b*). In this respect it should be noted that in addition to its impact on the overall growth and biomass of plants, when occurring during the reproductive phase of plant growth, a combination of drought and heat stress can have an even greater devastating impact on yield (Cohen *et al*., 2020). Thus, processes such as decreased fertilization, flower abortion, and production of fewer and smaller seeds have been reported to occur in different crops in response to a combination of drought and heat stress (Challinor *et al*., 2014; Lawas *et al*., 2018; Cohen *et al*., 2020). Deeper understanding of the effects of drought and heat stress combination on the reproductive processes of plants is therefore needed to develop crops with enhanced tolerance to this stress combination (Mittler, 2006; Mittler and Blumwald, 2010; Suzuki *et al*., 2014; Balfagón *et al*., 2020; Cohen *et al*., 2020; Zandalinas *et al*., 2020*b*).

In soybean (*Glycine max*), a considerable effort has been made to develop genotypes with enhanced yield production under conditions of heat or drought stress (*e.g*., Salem *et al*., 2007). However, how these genotypes perform under conditions of drought and heat stress combination is mostly unknown. Here, we studied the impact of water deficit and heat stress combination on the physiology and seed production (number of seeds and yield per plant) of two soybean genotypes, ‘Magellan’ and Plant Introduction (PI) 548313, which differ in their reproductive responses to conditions of heat stress. Our findings reveal that although seed number and yield per plant of PI 548313 were less affected by heat compared to Magellan, under conditions of water deficit and heat stress combination they decreased (similar to the effect of the stress combination on the Magellan genotype). Moreover, because the number of flowers and % pollen germination of PI 548313 remained high under heat stress or water deficit and heat stress combination, our findings suggest that the reduced seed numbers and yield per plant exhibited by PI 548313 under conditions of water deficit and heat stress combination could be a result of processes that occur at the stigma, ovaries and/or other parts of the flower following pollen germination.

## Materials and Methods

### Growth conditions

Seeds of ‘Magellan’ and PI 548313 were germinated in plastic trays filled with moist potting soil (ProMix, Premier Tech Horticulture, PA, USA) and inoculated with a commercial soybean inoculant (N-Dure, Verdesian, NC, USA). The trays were placed in growth chambers (BDR16, Conviron, Canada) at 24/20°C day/night temperature with a photoperiod of 12 h under a light intensity of 500 μmol photons m^−2^ s^−1^. After one week in the growth chamber, 48 uniform seedlings of each genotype were transferred into 5-L pots filled with exactly one kg of dry potting soil. Following transplanting, one liter of water was added to each pot containing one plant, and twelve plants per genotype were arranged randomly in each of four identical growth chambers (BDR16, Conviron, Canada) set to a photoperiod of 14 h at a light intensity of 1,000 μmol photons m^−2^ s^−1^ at the top of the canopy and day/night temperatures of 28/24°C. With the onset of reproductive development (appearance of the first flower, R1 developmental stage; Fehr *et al*., 1971), plants were divided into four treatments, Control (CT), water deficit (WD), heat stress (HS) and combined water deficit and heat stress (WD+HS), each assigned to one of the four growth chambers. Plants in the CT and HS treatment were well watered. In the WD and WD+HS treatments, plants were supplied with 30% of the water available for transpiration. To determine water amounts to be added, all pots were weighed daily. In the pots subjected to WD and WD+HS, the amount replenished brought the pot to a weight of about 1.6 kg while well-watered pots (CT and HS) were replenished to 2.4 kg. Day/night temperatures in CT and WD treatments were continued at 28/24°C day/night for 14/10 h, respectively. In order to simulate a peak of heat episode during mid-day, in the HS and WD+HS treatments, temperatures were ramped between 06:00-08:00 from 28 to 38°C, and maintained at 38°C for eight h (until 16:00). Then, ramping down to 28°C took place between 16:00-20:00. The night temperature in the heat treatment was maintained as in the CT treatment (24°C). Plants were maintained in these conditions for 20 days, at which point they were transferred to an environmentally controlled greenhouse at ~30/24°C, 60-70% RH and supplemental light from high pressure sodium bulbs of about 450 μmol photons m^−2^ s^−1^ at canopy, and scored for their seed production and yield per plant at physiological maturity as described below.

### Gas exchange and leaf temperature

Leaf gas exchange parameters (stomatal conductance, transpiration, photosynthesis and respiration) were determined at leaf temperatures of 28°C (CT and WD) or 38°C (HS and WD+HS), 1,500 μmol photons m^−2^ s^−1^ (10% blue light), 400 μmol CO_2_ mol^−1^ air and constant flow rate of 500 μmol m^−2^ s^−1^ using a portable photosynthesis system LICOR 6800 (LiCor, NE, USA), as recommended by the manufacturer, ten days into the stress period. For each treatment, six replicates were measured, and measurements took place between 10:00–12:00 using intact, fully expanded leaves exposed to direct light. Leaf temperature was determined with a handheld IR thermometer DX501-RS (Exergen, MA, USA), just prior to the gas exchange analysis.

### Reproductive traits

To quantify the reproductive response of soybean to the four treatments, the number of pods, the number of seeds in each pod, the number of nodes carrying pods, and the number of flowers per plant were manually counted during the entire course of the experiment. Individual seed weight was also determined using an analytical scale (Mettler Toledo, OH, USA), and yield per plant was measured as the total weight of all seeds produced by the plant at physiological maturity (growth stage R8; *i.e*., when 95 percent of the pods have obtained mature pod color; Fehr *et al*., 1971).

### Water relations

Relative water content (RWC) was based on fresh weight (FW), turgid weight (TW), and dry weight (DW) as follows: RWC (%) = (FW −DW)/(TW −DW) × 100. To determine FW, fully expanded leaves were harvested at 10:00 on day 16 of stress treatment initiation, weighed immediately and then placed into vials filled with distilled water for 24 h in the dark at 4°C. Leaf samples were then weighed to obtain TW. Samples were dried in an oven at 60°C for 3 days and weighed to determine DW. Midday leaf water potential (ψ) was measured on uppermost fully expanded trifoliate leaves from six plants per treatment with a pressure chamber (600 Pressure Chamber Instrument, PMS Instrument, OR, USA) at 10:00 on day 16 of stress treatment initiation. Immediately after excision, leaves were placed into the pressure chamber with the petiole distended from the chamber lid. The chamber was pressurized using a N2 tank, and Ψ was recorded upon the initial appearance of xylem sap on the cut petiole.

### Pollen germination assay

Pollen germination was measured according to (Djanaguiraman *et al*., 2019). Briefly, germination medium was prepared by dissolving 15 g sucrose, 0.01 g HBO_3_, 0.03 g CaNO_3_ and 0.5 g of agar in 100 ml of deionized water, heated until the agar completely dissolved, and dispensed onto microscopy slides to establish a layer for pollen germination. Pollen were collected from six different flowers located on the main stem per stress treatment between 10:00-12:00, 7-9 days into the stress treatments and placed on the pre-made microscope slides with germination medium under a dissecting scope (Leica S9i stereo microscope, Leica Microsystems, Wetzlar, Germany). The slides were placed in a square petri-dish and sealed with Parafilm (Bemis Company, WI, USA) for 30 min of incubation at 28°C or 38°C. The number of pollen grains with or without pollen tubes were counted under 40X magnification (EVOS XL, Invitrogen by Thermo Fisher Scientific, CA, USA) and percentage of germination was calculated. A total of at least 10 microscopic fields were counted per slide.

### RNA extraction

Ten days following stress initiation, young fully expanded Magellan leaves without visual tissue damage were harvested between 11:00-12:00, and immediately frozen in liquid nitrogen. Each sample was a pool of at least three different leaves from six different plants. Plant RNeasy kit (Qiagen, Germany) was used for RNA purification following the manufacturer’s instructions. Reverse transcription quantitative PCR (RT-qPCR) analysis was conducted as described in (Balfagón *et al*., 2019). Primer sequences used for the RT-qPCR analysis and their efficiency values are listed in Table S7.

### RNA-Seq data analysis

RNA sequencing was performed using NovaSeq 6000 at the University of Missouri DNA Core Facility as described in (Zandalinas *et al*., 2020*a*). RNA libraries were prepared for sequencing using standard Illumina protocols (Balfagón *et al*., 2019; Zandalinas *et al*., 2019, 2020*c*). Read quality control was performed using FastQC v1.20.0 (https://www.bioinformatics.babraham.ac.uk/projects/fastqc/). The reads were aligned to the reference genome of *Glycine max* v2.1 (downloaded from https://ftp.ncbi.nlm.nih.gov/genomes/all/GCF/000/004/515/GCF_000004515.5_Glycine_max_v2.1/), using STAR aligner v2.4.0.1 (Dobin *et al*., 2013). The corresponding genome annotation files were also downloaded from the same source. Differential gene expression analysis was performed using DESeq2 (Love *et al*., 2014). Differences in expression were quantified as the logarithm of the ratio of mean normalized counts between two conditions (log fold change). Differentially abundant transcripts were defined as those that have a log fold change with an adjusted p-value < 0.05 (negative binomial Wald test followed by a Benjamini-Hochberg correction; both integral to the DESeq2 package). Differentially expressed genes were classified into upregulated or downregulated based on significant positive or negative log fold change values, respectively. Venn diagram overlaps were calculated using VIB Bioinformatics and Evolutionary Genomics web tool (http://bioinformatics.psb.ugent.be/webtools/Venn/). Functional annotation and quantification of overrepresented GO terms (p-value < 0.05) were conducted using Panther (http://pantherdb.org/). Heat maps were generated using MeV v. 4.9.0 software (Saeed *et al*., 2006). Transcripts with significant changes in steady-state levels in response to the different treatments are shown in Tables S1-S6.

### Statistical analysis

Statistical analysis was performed using JMP 14 software (SAS Institute). Data was subjected to a two-way ANOVA, followed by Tukey–Kramer posthoc test (p < 0.05) when a significant difference was detected.

## Results

### Physiological characterization of soybean plants (cv. Magellan) subjected to a combination of water deficit and heat stress

To study the impact of water deficit (WD) and heat stress (HS) combination on the yield of soybean plants, we first established a set of environmental conditions for a WD+HS combination treatment (described in Materials and Methods) and determined its impact on the different physiological parameters of plants. As shown in Fig. 1, leaf temperature, RWC and leaf water potential of plants subjected to a combination of WD+HS were different from those of plants subjected to WD or HS alone. Leaves of plants subjected to the stress combination experienced a higher leaf temperature, a lower RWC and a lower leaf water potential, compared to leaves of plants subjected to WD or HS, constituting a unique combination of adverse conditions (Fig. 1). As shown in Fig. 2, the combination of WD+HS resulted in a significant decrease in net photosynthesis that was coupled with a decrease in transpiration and stomatal conductance and an increase in respiration. Interestingly, although stomatal conductance and transpiration of plants subjected to WD or WD+HS combination were similar, net photosynthesis of plants subjected to the stress combination was significantly lower than that of plants subjected to WD or HS (Fig. 2). These results suggest that in addition to the limitation in CO_2_ availability that results from the closure of stomata (decreased stomatal conductance), the higher leaf temperature and the lower water content of plants subjected to the stress combination (compared to WD or HS; Fig. 1) may cause a further reduction in photosynthesis in plants subjected to the stress combination.

**Fig. 1.**
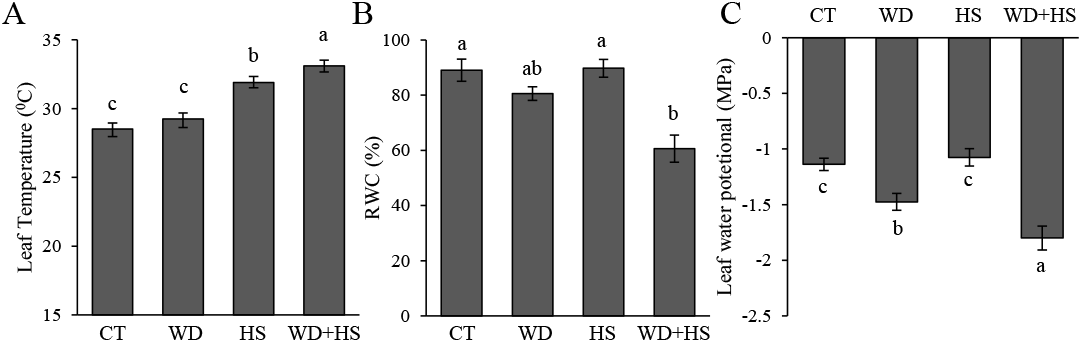
Leaf temperature, relative water content and leaf water potential of Magellan plants subjected to water deficit (WD), heat stress (HS) and the combination of water deficit and heat stress (WD + HS). (A) Leaf temperature, (B) relative water content and (C) leaf water potential. ANOVA, SD, N=6, different letters denote statistical significance at P < 0.05. *Abbreviations used*: CT, control; water deficit, WD; heat stress, HS; combination of water deficit and heat stress, WD + HS; T, temperature; RWC, relative water content.

**Fig. 2.**
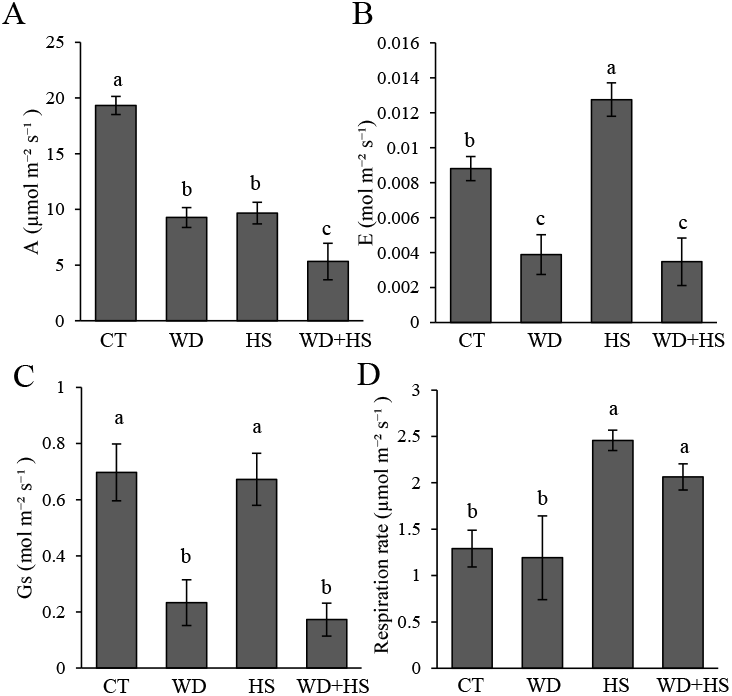
Physiological parameters of Magellan plants subjected to water deficit (WD), heat stress (HS) and the combination of water deficit and heat stress (WD + HS). (A) Photosynthetic rate, (B) transpiration, (C) stomatal conductance, and (D) respiration. ANOVA, SD, N=6, different letters denote statistical significance at P < 0.05. *Abbreviations used*: CT, control; water deficit, WD; heat stress, HS; combination of water deficit and heat stress, WD + HS.

### RNA-Seq analysis of soybean cv. Magellan subjected to a combination of WD+HS

To further study the response of soybean plants subjected to a combination of WD+HS and to identify different transcripts unique to this state of stress combination, we conducted an RNA-Seq analysis of leaves subjected to the stress combination shown in Figs. 1 and 2. This analysis revealed that the steady-state levels of 5,188 and 5,224 transcripts was enhanced or decreased, respectively, in plants subjected to a combination of WD+HS (Figs. 3, S1, and S2). Of these, the steady-state levels of 2,584 and 3,103 transcripts was specifically altered (enhanced or decreased, respectively) in response to the stress combination (Fig. 3). As with previous transcriptomics studies of drought and heat stress combination in Arabidopsis (Rizhsky *et al*., 2004; Rasmussen *et al*., 2013; Prasch and Sonnewald, 2013; Shaar-Moshe *et al*., 2017; Zandalinas *et al*., 2020*b*), the degree of overlap between the transcriptomic responses to WD or HS was low, with only 202 and 190 transcripts commonly up and down regulated, respectively, between these two different stresses (out of 2,084 and 1,536 transcripts up or down regulated, respectively, in response to WD, and 3,563 and 2,890 transcripts up and downregulated, respectively, in response to HS; Fig. 3). These findings highlight the distinctiveness of plant responses to a combination of WD+HS, two stresses that have a relatively low similarity in terms of transcriptomic responses. In line with this low similarity, the steady-state level of only 176 and 166 transcripts was up- or downregulated, respectively, in response to all treatments (WD, HS and WD+HS combination; Fig. 3). As also shown in Fig. 3, the groups of transcripts specifically upregulated in response to WD (1,194), HS (1,621), WD+HS (2,584), or common to all stresses (176) were enriched in transcripts involved in transferase, hydrolase and oxidoreductase activity. Functional terms associated with biological processes that were enriched in the sets of upregulated transcripts specific to WD, HS, WD and HS, or common to all stresses, included response to stimulus, signal transduction, cell communication and oxidation-reduction. In contrast, the sets of downregulated transcripts representing these same treatment groups were not enriched in “response to stimulus” transcripts (Fig. 3).

**Fig. 3.**
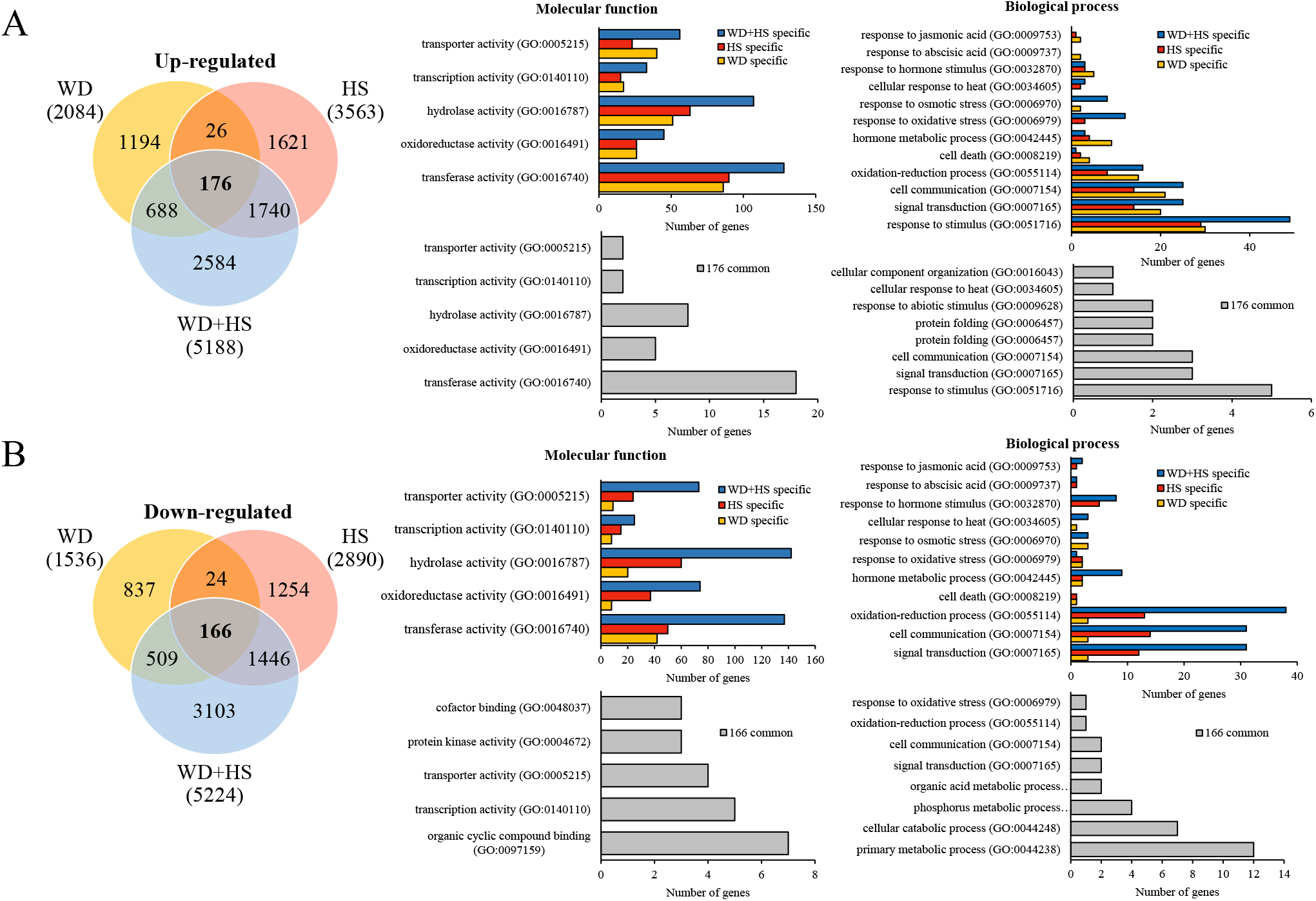
Transcriptomics analysis of Magellan plants subjected to water deficit (WD), heat stress (HS) and the combination of water deficit and heat stress (WD + HS). (A) Venn diagram depicting the overlap between transcripts upregulated in response to WD, HS and WD + HS (left) and bar graphs of biological process and molecular function (GO) annotations for upregulated transcripts specific to WD, specific to HS, specific to WD + HS, and common among all stress conditions (right). (B) Venn diagram depicting the overlap between transcripts downregulated in response to WD, HS and WD + HS (left) and bar graphs of biological process and molecular function (GO) annotations for downregulated transcripts specific to WD, specific to HS, specific to WD + HS, and common among all stress conditions (right). *Abbreviations used*: CT, control; water deficit, WD; heat stress, HS; combination of water deficit and heat stress, WD + HS.

To identify regulatory proteins that could potentially be used in future breeding or genetic transformation studies to enhance the tolerance of soybean plants to a combination of WD+HS, we conducted an expression analysis of different transcription factor (TF) genes during WD, HS, and WD+HS combination. As shown in Fig. 4, many transcripts belonging to the heat shock transcription factor (HSF), MYB, WRKY and dehydration responsive element binding (DREB) families, were altered in their expression in response to the different treatments. However, only a selected few (MYBs 84, 108 and 184, HSF1, and ERF060-like) were upregulated in response to all 3 treatments (Fig. 4). Downregulated TFs demonstrated a similar behavior with only 5 (ABR1, ERF-like, EREB, WRKY1 and WRKY3) suppressed in response to all treatments (Fig. 4). The findings presented in Fig. 4 identify several transcriptional regulators that could be used in future efforts to enhance the tolerance of soybean plants to a combination of WD+HS. However, they also demonstrate that a combinatorial, rather than an additive, approach to TF network regulation might be needed to enhance tolerance to stress combination, as proposed for Arabidopsis by Zandalinas *et al*., (2020*b*).

**Fig. 4.**
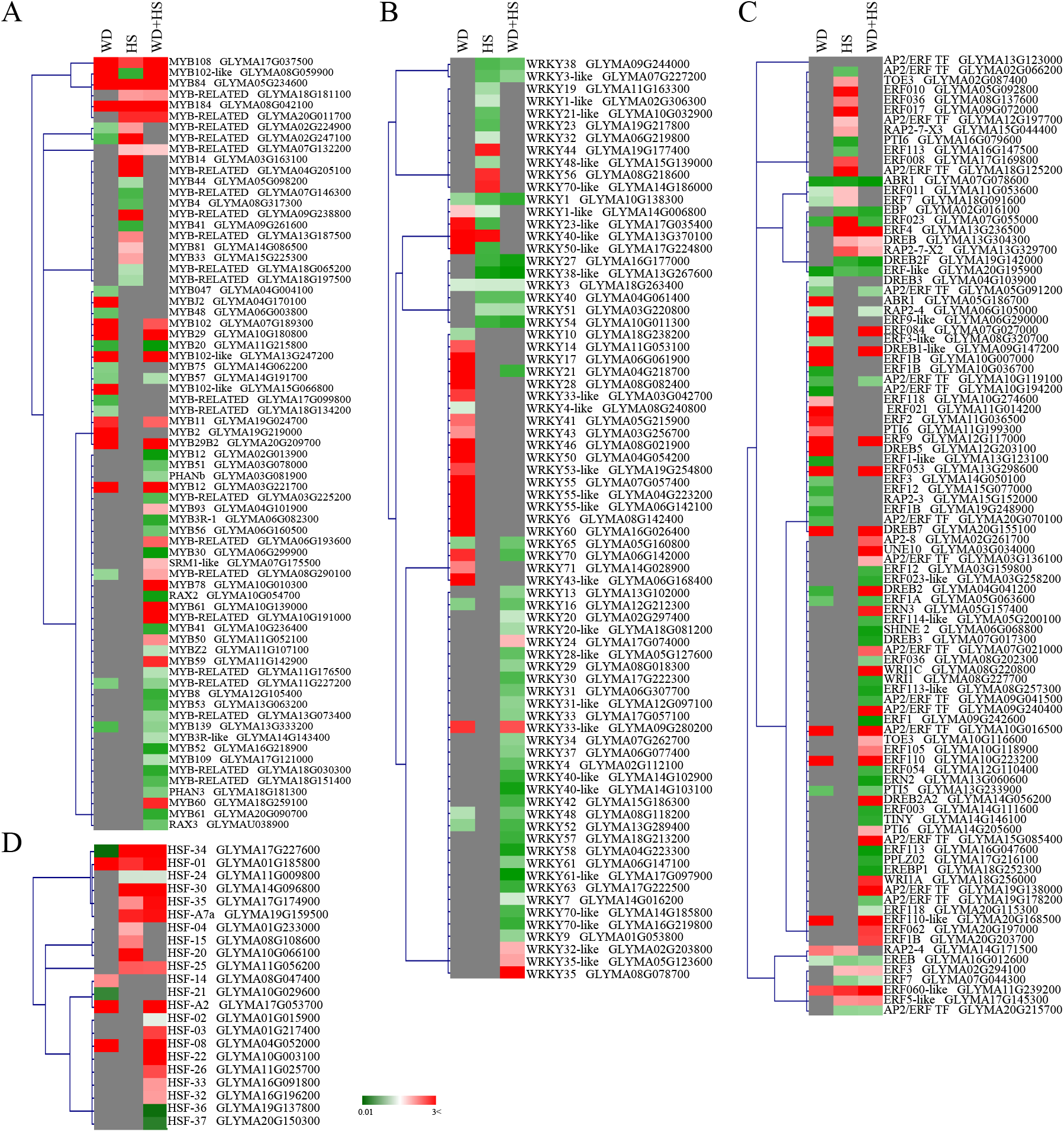
Expression of selected transcription factor families in Magellan plants subjected to water deficit (WD), heat stress (HS) and the combination of water deficit and heat stress (WD + HS). Heat maps showing the expression level of (A) MYBs, (B) WRKYs, (C) AP2-ERFs, and (D) HSFs, in Magellan plants subjected to the different stresses. *Abbreviations used*: CT, control; water deficit, WD; heat stress, HS; combination of water deficit and heat stress, WD + HS.

### Physiological characterization of soybean PI 548313 subjected to a combination of WD+HS

To determine whether enhanced tolerance to HS in soybean would convey higher yield under conditions of WD+HS combination, we obtained a soybean genotype that exhibited in preliminary studies an above average relative number of seeds per plant under HS compared to ambient temperature (Plant Introduction, PI 548313; FBF, personal communication). To compare PI 548313 to our control Magellan line, we subjected it to the same WD+HS combination treatment used for Magellan (Figs. 1–4) and determined its physiological responses. As shown in Fig. 5, net photosynthesis, transpiration and stomatal conductance of PI 548313 responded in a similar manner to that of Magellan to the different stress conditions (Fig. 2). In contrast, compared to Magellan, PI 548313 displayed a relatively lower leaf water potential during HS (Figs. 1 and 5).

**Fig. 5.**
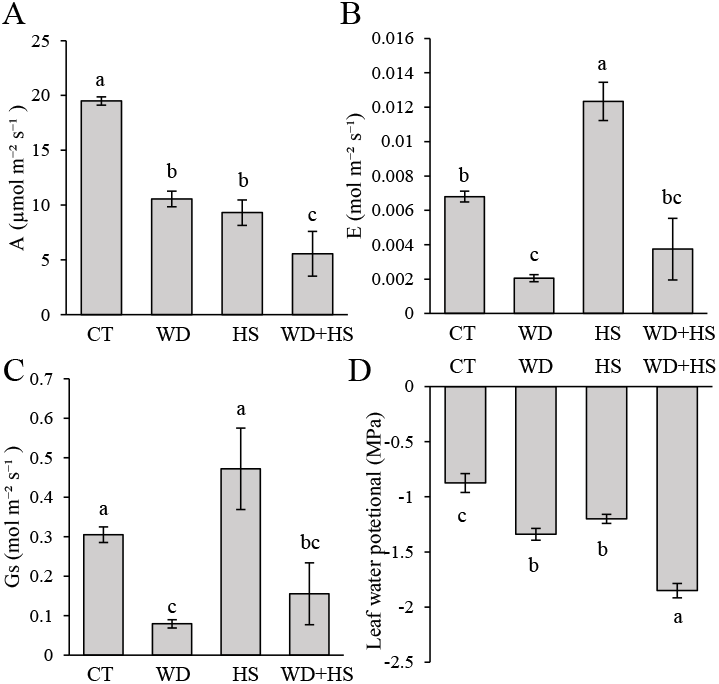
Physical and physiological parameters of PI 548313 plants subjected to water deficit (WD), heat stress (HS) and the combination of water deficit and heat stress (WD + HS). (A) Photosynthetic rate, (B) transpiration, (C) stomatal conductance, and (D) leaf water potential, of PI 548313 plants subjected to the different stresses. ANOVA, SD, N=6, different letters denote statistical significance at P < 0.05. *Abbreviations used*: CT, control; water deficit, WD; heat stress, HS; combination of water deficit and heat stress, WD + HS.

### Seed production and pollen germination of Magellan and PI 548313 under conditions of WD+HS combination

To address the question of whether the ability of PI 548313 to produce above average relative number of seeds per plant under HS will enable this genotype to produce more seeds per plant under conditions of WD+HS combination, we grew the two genotypes side-by-side under the conditions established in Figs. 1, 2, and 5, and compared their total number of seeds per plant, yield per plant, and number of flowers and pods per plant under the different stress treatments. As shown in Fig. 6A, total seed number per plant of Magellan decreased under WD, HS and WD+HS combination, with the stress combination being the most severe, followed by HS and then WD. Compared to Magellan, PI 548313 displayed a similar decrease in the number of seeds produced per plant in response to WD but had a significantly lower decrease in seed numbers in response to HS. Interestingly, in response to the WD+HS combination both genotypes displayed a further decrease in the number of seeds produced per plant compared to WD or HS (Fig. 6A). A similar impact of WD, HS and WD+HS was observed on yield per plant (Fig. 6B). These findings indicate that although the impact of heat on PI 548313 was not as severe as that on Magellan, in response to the stress combination both the PI 548313 and Magellan suffered a further reduction in number of seeds and yield per plant. Interestingly, in response to HS both genotypes displayed a decrease in average seed weight (Fig. S3) that was not observed in WD or WD+HS combination.

**Fig. 6.**
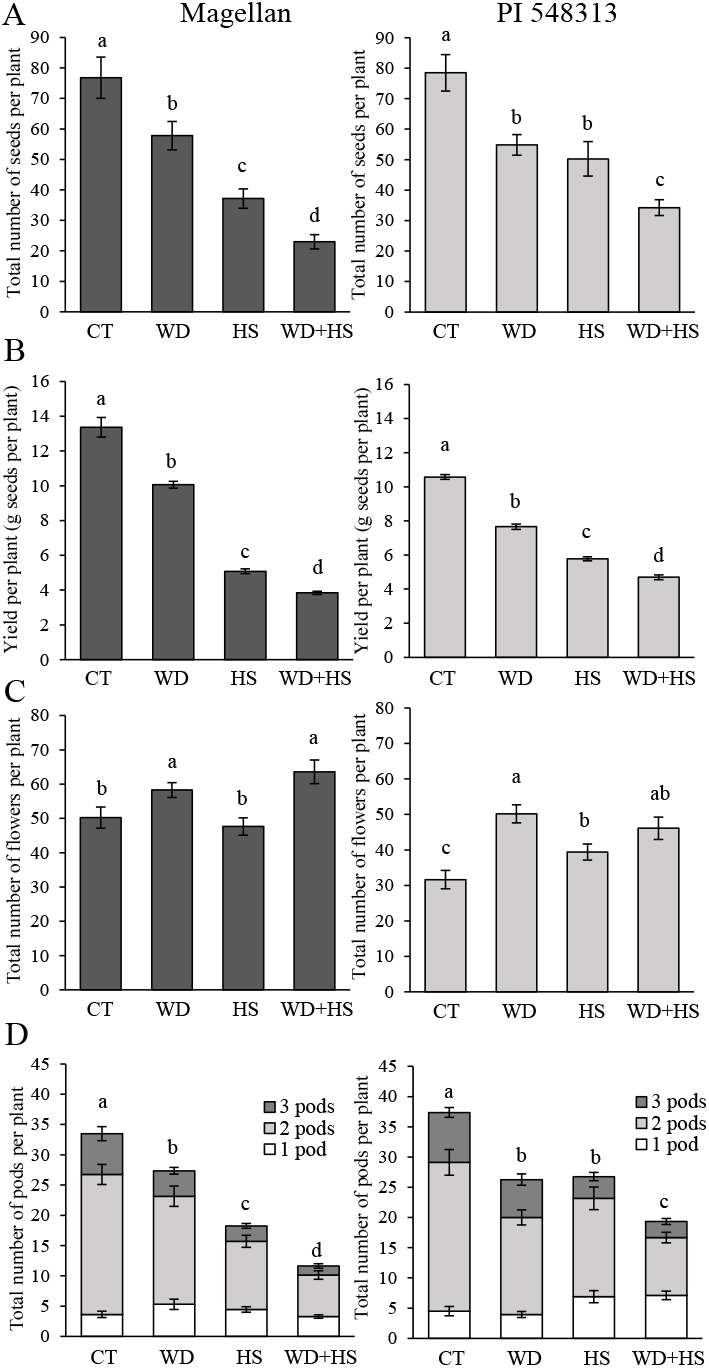
Seed, flower and pod production of Magellan and PI 548313 plants subjected to water deficit (WD), heat stress (HS) and the combination of water deficit and heat stress (WD + HS). (A) Total number of seeds per plant, (B) Yield per plant (g seeds per plant), (C) Total number of flowers per plant, and (D) Total number of pods per plant of Magellan (left) and PI 548313 (right) plants subjected to the different stresses were determined during the course of the stress treatments and at physiological maturity. ANOVA, SD, N=6, different letters denote statistical significance at P < 0.05. *Abbreviations used*: CT, control; water deficit, WD; heat stress, HS; combination of water deficit and heat stress, WD + HS.

As shown in Fig. 6C, the decrease in total seed production per plant observed in both Magellan and PI 548313 under the stress combination (Fig. 6A) was not the result of a decrease in the number of flowers produced per plants. In fact, both Magellan and PI 548313 produced a high number of flowers in response to the stress combination. These findings suggest that the Magellan and PI 548313 genotypes growing under the stress combination were not acutely deficient in energy and/or resources needed to produce a sufficient number of flowers (that could have explained the decrease in total seeds produced per plant). The lower number of seeds produced by the Magellan and PI 548313 genotypes under the stress combination may therefore result from processes such as decrease in pollen viability, pollen tube growth, unsuccessful fertilization, inhibited embryogenesis, and/or enhanced abortion of pods and/or seed.

As shown in Fig. 6D, the high number of flowers produced in both Magellan and PI 548313 in response to the stress combination (Fig. 6C) did not result in a higher number of pods per plant at physiological maturity. In fact, the number of pods in both Magellan and PI 548313 corresponded very well to the total number of seeds per plant (Fig. 6A). The number of pods per plant in Magellan decreased under WD, HS and WD+HS combination, with the stress combination being the most severe, followed by HS and then WD. By contrast, while the number of pods per plant in PI 548313 displayed a similar decrease in response to WD, PI 548313 had a significantly higher number of pods per plant compared to Magellan in response to HS (Fig. 6D). The results presented in Fig. 6D also demonstrate that with the reduced number of pods per plant, the number of pods with 2 or 3 seeds per pod was also reduced in response to the different stress conditions, whereas the number of pods with 1 seed per pod was relatively stable, or even increased in PI 548313 in response to HS or the WD+HS stress combination. The results presented in Fig. 6C and 6D suggest that although large numbers of flowers were produced per plant during a combination of WD+HS, the success rate of fertilization, embryogenesis and/or other post fertilization processes was decreased under the stress combination.

To test whether the decrease in total seed and pod number per plant in response to the stress combination (Fig. 6A and 6D) was a result of a decrease in pollen viability caused by the stress combination, we measured % pollen germination of the two genotypes. As shown in Fig. 7, pollen viability (as evident by pollen germination rate) was decreased in the two genotypes in response to WD and more severely in response to HS. Nevertheless, the % germination of pollen obtained from plants subjected to a combination of WD+HS was not significantly different from that of pollen obtained from plants subjected to HS. This finding suggests that the decrease in yield caused by the WD+HS combination might be a result of the stress combination interfering with reproductive processes other than those determining pollen germination rates.

**Fig. 7.**
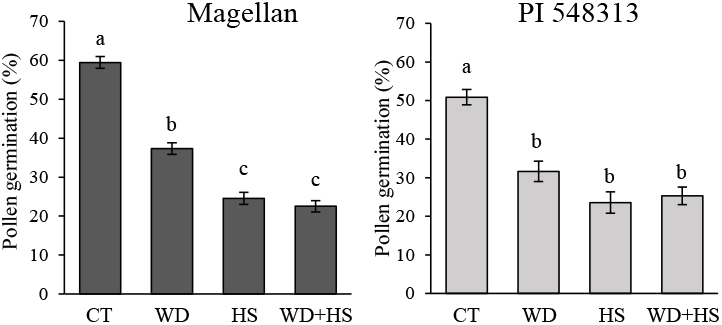
Germination of pollen collected from Magellan and PI 548313 plants subjected to water deficit (WD), heat stress (HS) and the combination of water deficit and heat stress (WD + HS). ANOVA, SD, N=6, different letters denote statistical significance at P < 0.05. *Abbreviations used*: CT, control; water deficit, WD; heat stress, HS; combination of water deficit and heat stress, WD + HS.

## Discussion

A combination of WD+HS can occur during the vegetative, reproductive or both phases of plant growth with different impacts on the biomass and yield of different crops (Cohen *et al*., 2020; Zandalinas *et al*., 2020*b*). During the vegetative growth phase of plants, the stress combination can impact photosynthesis, carbon assimilation and plant growth, while during the reproductive growth phase it can additionally impact flowering, fertilization, and seed production (Mittler and Blumwald, 2010; Balfagón *et al*., 2020; Cohen *et al*., 2020). In our study we compared the response of two different soybean genotypes to a combination of WD+HS occurring during their reproductive growth phase. The two genotypes we compared differed in their ability to produce seeds under conditions of HS (Fig. 6). Interestingly, the physiological responses of the two genotypes to the different stress treatments were very similar (Figs. 1, 2, and 5), suggesting that the advantage of PI 548313 over Magellan in seed production per plant during HS (Fig. 6A) was not a result of differences between these two genotypes in tolerance to HS of leaf-level physiological processes. In support of this possibility was also the ability of both genotypes to produce a similar or higher number of flowers under the different stress conditions compared to control (Fig. 6C). Compared to control conditions, the PI 548313 genotype was however able to produce more flowers in response to HS than Magellan (Fig. 6C). The PI 548313 genotype might therefore be different from Magellan in its ability to generate flowers and complete its reproductive process (higher pod number and total seed number per plant) under conditions of HS (Fig. 6). Interestingly, although PI 548313 was superior to Magellan in seed production per plant during HS, this superiority did not prevent a further decrease in the number of seeds or yield per plant once HS was combined with WD (WD+HS; Fig. 6). Because the two genotypes displayed a similar physiological response to the different stress treatments (Figs. 2 and 5), the impact of the stress combination on seed production in both genotypes appears to be a specific effect of the stress combination on reproductive tissues. Additional studies are of course needed to carefully measure and compare biomass and yield between the two genotypes under the different conditions reported here, applied and extended upon to field conditions.

Many studies have linked the decrease in yield occurring during HS with a decrease in pollen viability (*e.g*., Kumar *et al*., 2015; Lawas *et al*., 2018; Cohen *et al*., 2020). Indeed, pollen germination of both Magellan and PI 548313 decreased in response to WD, HS, or WD+HS (Fig. 7). Interestingly, the pollen germination rates of both the PI 548313 and Magellan genotypes were similar under conditions of HS and WD+HS (Fig. 7). Despite this similarity, seed number and yield per plant of both Magellan and PI 548313 suffered a similar further decrease in response to the combination of WD+HS (Fig. 6). The similar pollen germination rates observed between HS and WD+HS in both genotypes (Fig. 7), was therefore not sufficient to maintain similar seed number or yield per plant in response to the stress combination (Fig. 6). Taking into account that the number of flowers in both genotypes increased during WD+HS combination (Fig. 6), and that pollen germination rates of both genotypes were similar under conditions of HS and WD+HS combination (Fig. 7), our findings suggest that at least when it comes to Magellan and PI 548313, the cause of yield reduction under conditions of WD+HS combination appears to be a result of disruption in processes that occur downstream to pollen germination on the flower pistil. Thus, processes such as pollen tube growth, ovule and egg development, fertilization and/or embryogenesis might be adversely affected under conditions of WD+HS combination in the two genotypes. Further analysis of these processes is therefore needed to clearly identify the cause of yield reduction during a combination of WD+HS in Magellan and PI 548313.

Our analysis of the physiological response of soybean to a combination of WD+HS revealed that the conflicting stomatal response that occurs in Arabidopsis and tobacco in response to this stress combination (Rizhsky *et al*., 2002, 2004) also occurs in soybean (Fig. 2). Thus, during HS stomata open, during WD they close, and during a combination of WD+HS they remain closed. Similar to Arabidopsis and tobacco, the overall outcome of this conflict is that the leaf temperature of plants subjected to the stress combination is higher, by about 1-2°C, compared to that of plants subjected to heat alone (due to the inability of plants subjected to the stress combination to cool their leaves by transpiration; Fig. 2; Rizhsky *et al*., 2002, 2004; Mittler, 2006). The higher leaf temperature of plants subjected to the stress combination could further explain the inhibition of photosynthesis that was higher in plants subjected to WD+HS combination compared to WD alone (Fig. 2), even though stomatal conductance of plants subjected to WD or a combination of WD+HS was similar (Fig. 2). The negative impact of stress combination on photosynthesis was recently reported for a combination of excess light and HS in Arabidopsis (Balfagón *et al*., 2019), and it is possible that a similar impact on D1 turnover and photosynthesis occurs in soybean under conditions of WD+HS combination. Further studies are of course needed to determine the cause of photosynthesis reduction under conditions of stress combination in soybean. Taken together, it appears however that although Arabidopsis is not a good experimental system to study the impact of WD+HS combination on yield (Vile *et al*., 2012; Cohen *et al*., 2020), it might be a good system to study other effects of this stress combination on vegetative tissues of plants (Rizhsky *et al*., 2004; Rasmussen *et al*., 2013; Prasch and Sonnewald, 2013; Choudhury *et al*., 2017; Zandalinas *et al*., 2020*a*,*b*; Balfagón *et al*., 2020).

Our RNA-Seq analysis of the response of soybean to a combination of WD+HS demonstrated the uniqueness of this response, as well as identified thousands of different transcripts that are exclusively expressed (induced or suppressed) during a combination of WD+HS (Figs. 3 and 4). Furthermore, our analysis identified a number of different TFs that are induced under all stress conditions studied (WD, HS and WD+HS combination; Figs. 3 and 4). These could be used in future breeding and genetic engineering efforts to improve the tolerance of soybean to conditions of stress combination (Mittler and Blumwald, 2010; Bailey-Serres *et al*., 2019). Although it is possible that overexpressing one or more of these TFs will improve the tolerance of soybean to WD+HS stress combination, it is more likely that tolerance to stress combination would require changes in the expression of more than a single regulatory gene, and that multiple loci are involved in the response of plants to stress combination (Mittler and Blumwald, 2010; Yuan *et al*., 2019; Shaar-Moshe *et al*., 2019; Zandalinas *et al*., 2020*b*) Additionally, it is possible that a combination of different TFs expressed in the correct order in specific tissues might be needed (a combinatorial response; Zandalinas *et al*., 2020*b*) Future studies are therefore needed to address these possibilities as well as to decipher the different pathways involved in the response of soybean to stress combination. These should contain more time points and focus on vegetative as well as reproductive tissues. In addition, more careful attention is needed to the manner in which the stress combination is applied to plants. For example, polyethylene glycol should not be used to induce WD conditions in soybean plants grown in soil (Wang *et al*., 2018) since it causes responses that do not mimic WD under field conditions (Mittler and Blumwald, 2010). The RNA-Seq data sets reported in our study can serve as an initial baseline and reference source for the response of soybean to a combination of WD+HS, nevertheless they should be followed in future studies by more detailed molecular analyses of reproductive tissues, that appear to play a much more important role in yield penalty under the stress combination than effects on leaf tissues (Figs. 6 and 7).

## Supplementary data

**Fig. S1.**
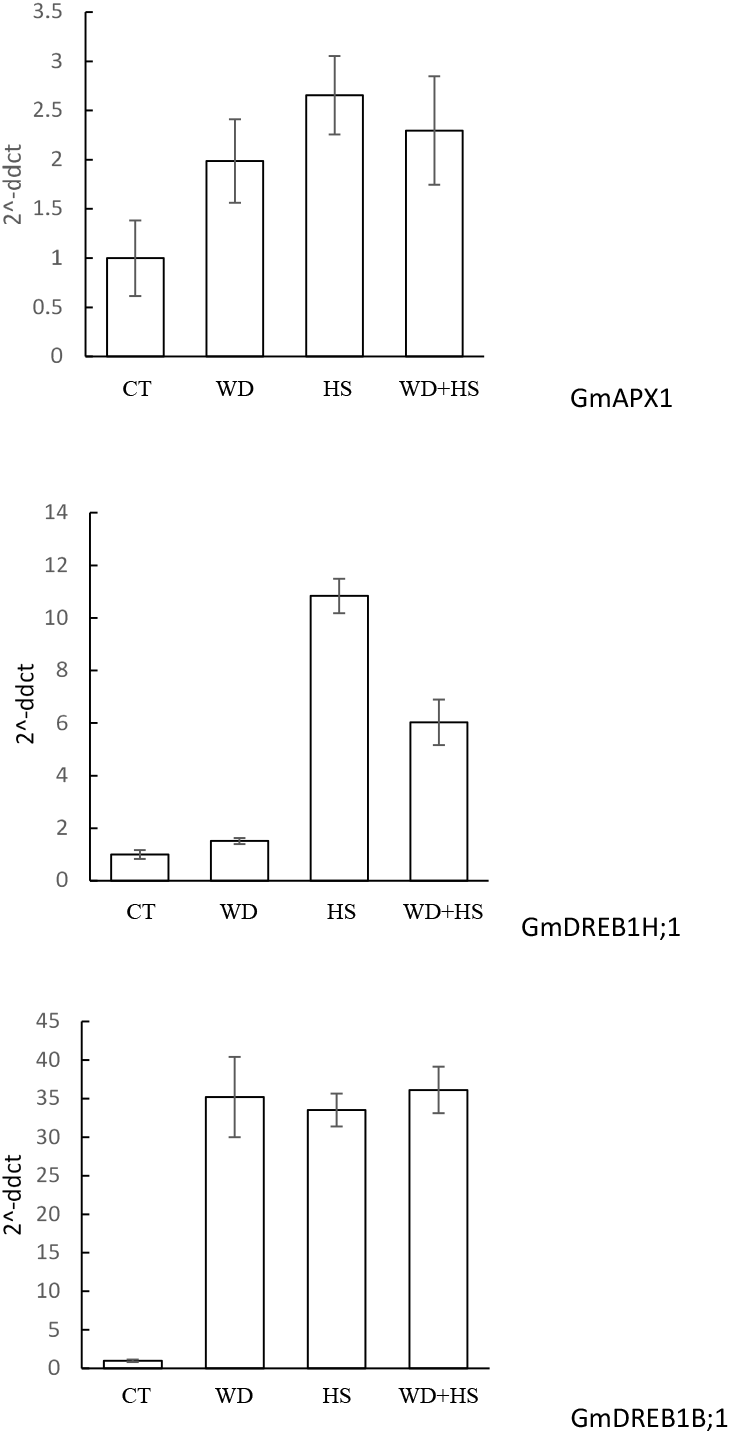
RT-qPCR analysis of selected stress-response transcripts in the RNA samples used for RNA-Seq analysis.

**Fig. S2.**
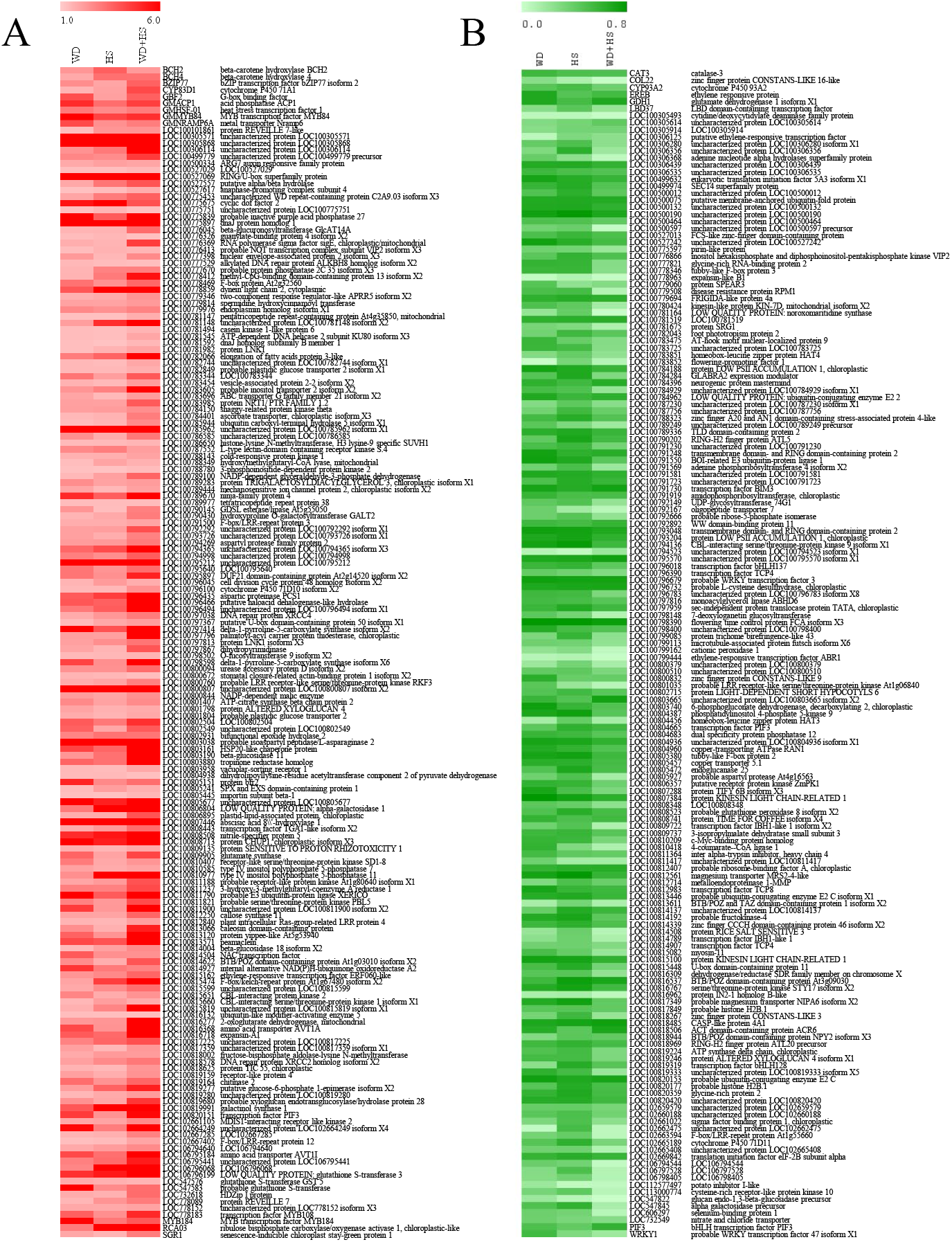
A heat map showing the expression pattern of 176 (A) and 166 (B) transcripts up or downregulated, respectively, in response to all stress conditions.

**Fig. S3.**
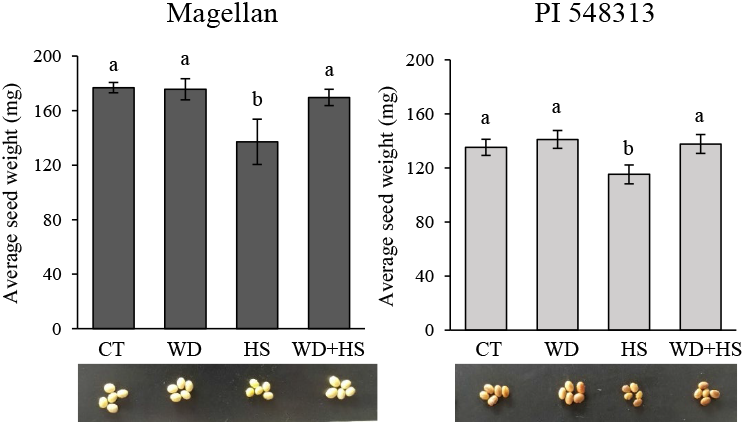
Individual seed weight of Magellan (Left) and PI 548313 (Right) as affected by heat (HS), water deficit (WD) and their combination (WD+HS). Results are shown as average and SE different of 12 plants per treatment, sampled at day 20 of stress imposition. Bars labeled with different letters are significantly different (P ≤ 0.05) according to one way ANOVA followed by Tukey’s test.

**Table S1.** Transcripts significantly upregulated compared to control (P < 0.05) in Magellan subjected to water deficit.

**Table S2.** Transcripts significantly upregulated compared to control (P < 0.05) in Magellan subjected to heat stress.

**Table S3.** Transcripts significantly upregulated compared to control (P < 0.05) in Magellan subjected to the combination of water deficit and heat stress.

**Table S4.** Transcripts significantly downregulated compared to control (P < 0.05) in Magellan subjected to water deficit.

**Table S5.** Transcripts significantly downregulated compared to control (P < 0.05) in Magellan subjected to heat stress.

**Table S6.** Transcripts significantly downregulated compared to control (P < 0.05) in Magellan subjected to the combination of water deficit and heat stress.

**Table S7.** Primers used for RT-qPCR

## Acknowledgements

The RNA sequencing service was provided by the University of Missouri DNA Core Facility. This work was supported by funding from the National Science Foundation (NSF-BSF MCB-1936590, IOS-1932639, and IOS-1353886), and the University of Missouri.

## Author contribution

I.C., S.I.Z, S.S. and Y.F. performed experiments and analyzed the data. R.M., F.B.F, I.C. and S.I.Z. designed experiments and analyzed the data. R.K.A. coordinated bioinformatics analysis. S.I.Z., F.B.F., R.K.A., I.C. and R.M. wrote the manuscript. All authors read and approved the manuscript.

## Data availability statement

The RNA-seq data analyzed in this study were deposited in the Gene Expression Omnibus (GEO) database, https://www.ncbi.nlm.nih.gov/geo, under the accession no. GSE153951.

